# Biomedical Text Mining for Research Rigor and Integrity: Tasks, Challenges, Directions

**DOI:** 10.1101/108480

**Authors:** Halil Kilicoglu

## Abstract

An estimated quarter of a trillion US dollars is invested in the biomedical research enterprise annually. There is growing alarm that a significant portion of this investment is wasted, due to problems in reproducibility of research findings and in the rigor and integrity of research conduct and reporting. Recent years have seen a flurry of activities focusing on standardization and guideline development to enhance the reproducibility and rigor of biomedical research. Research activity is primarily communicated via textual artifacts, ranging from grant applications to journal publications. These artifacts can be both the source and the end result of practices leading to research waste. For example, an article may describe a poorly designed experiment, or the authors may reach conclusions not supported by the evidence presented. In this article, we pose the question of whether biomedical text mining techniques can assist the stakeholders in the biomedical research enterprise in doing their part towards enhancing research integrity and rigor. In particular, we identify four key areas in which text mining techniques can make a significant contribution: plagiarism/fraud detection, ensuring adherence to reporting guidelines, managing information overload, and accurate citation/enhanced bibliometrics. We review the existing methods and tools for specific tasks, if they exist, or discuss relevant research that can provide guidance for future work. With the exponential increase in biomedical research output and the ability of text mining approaches to perform automatic tasks at large scale, we propose that such approaches can add checks and balances that promote responsible research practices and can provide significant benefits for the biomedical research enterprise.

**Supplementary information:** Supplementary material is available at *BioRxiv*.

## 1 Introduction

Lack of reproducibility and rigor in published research, a phenomenon sometimes referred to as the “reproducibility crisis”, is a growing concern in science. In a recent Nature survey, 90% of the responding scientists agreed that there was a crisis in science (Baker, 2016a). It has become routine in recent years for scientific journals as well as for news media to publish articles discussing various aspects of this crisis as well as proposals and initiatives to address them. The reproducibility problem is perhaps most acutely felt in biomedical research, where the stakes are high due to the size of research investment and impact on public health. In 2010, the global spending on research in life sciences (including biomedical) was US$240 billion (Røttingen *et al.*, 2013). The problems in reproducibility and rigor of published research means that a portion of this expenditure is wasted. Chalmers and Glasziou (2009) estimate that avoidable waste accounts for approximately 85% of research investment.

A variety of factors, occurring at various stages of research and attributable to different stakeholders, can lead to reproducibility issues and, ultimately, waste. For example, at the conception, the scientist, unaware of the published literature, may propose to address a research question that can already be answered with existing evidence, or may fail to select the appropriate experimental design and methods (Chalmers and Glasziou, 2009; Collins and Tabak, 2014). As the research is being conducted, the investigator, overwhelmed with administrative tasks, may not be able to provide adequate training/supervision to lab staff (Barham *et al.*, 2014), who do not validate their experiments sufficiently. Only a subset of data that yields statistical significant results may be reported, while negative results may be discarded completely (*p*-hacking, *selective reporting*, or *publication bias*) (Head *et al.*, 2015; Dwan *et al.*, 2014; Chan *et al.*, 2004). The authors may fail to identify the model organisms, antibodies, reagents necessary for other researchers to replicate the experiments (Vasilevsky *et al.*, 2013). Journal editors, valuing novelty over reproducibility, may be reluctant to publish negative results or replication studies (Collins and Tabak, 2014). A peer reviewer may miss methodological problems with the manuscript. An institutional review board (IRB) may fail to follow up on biased underreporting of the research that they approve (Chalmers, 2002). Funding agencies may put too much emphasis on number of publications, citation counts, and research published in journals with high impact factors for rewarding research grants (Collins and Tabak, 2014; Bowen and Casadevall, 2015). The so-called “publish or perish” culture at academic institutions can create pressure to maximize research quantity with diminishing quality (Collins and Tabak, 2014).

While research rigor and reproducibility in biomedical research is not a recent problem, discussions of the “reproducibility crisis” are largely due to several recent high-profile studies. In one of the pioneering studies, Ioannidis (2005) demonstrated how reliance on hypothesis testing in biomedical research frequently results in false positive results, which he attributed to a variety of factors, such as effect size, flexibility in study design, and financial interest and prejudice. More recently, Begley and Ellis (2012) were unable to reproduce the findings reported in 47 of 53 landmark hematology and oncology studies. Studies with similar findings were conducted in other fields, as well (Kyzas *et al.*, 2007; Prinz *et al.*, 2011; Open Science Collaboration, 2015). Lack of reproducibility and rigor can mostly be attributed to questionable research practices (honest errors, methodological problems). At the extreme end of the reproducibility spectrum, fraudulent science and retractions constitute a small but growing percentage of the published literature. The percentage of retracted articles in PubMed has increased about 10-fold since 1975 and 67.4% are attributable to scientific misconduct: fraud, duplicate publication, and plagiarism (Fang *et al.*, 2012). Due to their pervasiveness, however, questionable research practices can be much more detrimental to science (Bouter *et al.*, 2016). Biomedical research outcomes, estimated by life expectancy and novel therapeutics, have remained constant despite rising investment and scientific knowledge in the last five decades, partly attributed to non-reproducibility (Bowen and Casadevall, 2015). Such evidence establishes the lack of reproducibility and rigor as a major problem that can potentially undermine trust in biomedical research enterprise. All stakeholders involved in the biomedical research enterprise have a responsibility to ensure the accuracy, verifiability, and honesty of research conduct and reporting to reduce waste and increase value.

Towards increased rigor and reproducibility, initiatives focusing on various aspects of reproducible science have been formed and they have been publishing standards, guidelines, and principles. These include ICMJE trial registration requirement (De Angelis *et al.*, 2004), NIH efforts in enhancing research reproducibility and transparency (Collins and Tabak, 2014) and data discovery (Ohno-Machado *et al.*, 2015), in addition to reporting guidelines (Simera *et al.*, 2010; Nosek *et al.*, 2015), data sharing principles (Wilkinson *et al.*, 2016), conferences (e.g., World Conference on Research Integrity), journals (e.g., Research Integrity and Peer Review), and centers (e.g., Center for Open Science, METRIC) dedicated to these topics.

We have used several terms (reproducibility, replication, rigor, integrity) somewhat interchangeably to describe related phenomena that differ in some fundamental aspects. The semantics of these terms are still somewhat muddled, leading to confusion and potentially hampering efforts to fix the problems (Baker, 2016b). To better focus the remainder of this paper, we use the definitions below, provided in Bollen *et al.* (2015).

- **Reproducibility:** The ability to duplicate the results of a prior study using the same materials and procedures as were used by the original investigator.
- **Replicability:** The ability to duplicate the results of a prior study if the same procedures are followed but new data are collected.
- **Generalizability:** Whether the results of a study apply in other contexts or populations that differ from the original one (also referred to as *translatability*).

Results that are reproducible, replicable, and generalizable are referred to as being *robust*.

The notions of rigor and integrity are also invoked to discuss related phenomena. The definitions below are taken from NIH’s Grants and Funding resources:

- **Rigor:** Strict application of the scientific method to ensure robust and unbiased experimental design, methodology, analysis, interpretation and reporting of results^1^.
- **Integrity:** The use of honest and verifiable methods in proposing, performing, and evaluating research, reporting results with attention to adherence to rules, regulations, guidelines, and following commonly accepted professional codes and norms^2^.

While reproducibility is often used as an umbrella term to cover all these related issues, in the remainder of this paper, we use it in the limited sense given above. Instead, we focus on the notions of research rigor and integrity, because i) these notions emphasize reporting over duplication of prior experiments, and ii) we are mainly interested in whether/how mining of textual research artifacts can contribute to openness and transparency in biomedical science.

Mining of textual biomedical research artifacts is in the purview of biomedical natural language processing (referred to as bioNLP henceforth), a field at the intersection of natural language processing (NLP) and biomedical informatics. In this article, we assume basic knowledge of bioNLP; for introductions and recent surveys, see Ananiadou and McNaught (2006); Cohen and Demner-Fushman (2014); Gonzalez *et al.* (2016). In the next section, we turn to our original question: can bioNLP provide tools that can help stakeholders in enhancing rigor and integrity of biomedical research?

## BioNLP for Research Rigor and Integrity

Text mining is largely concerned with unstructured text, the primary means of communication for biomedical research. Unstructured biomedical text comes in various forms of textual artifacts, including:

- *Proposals* (grant applications, protocols) authored by scientists and assessed by funding agencies, reviewers, and IRBs
- *Manuscript submissions*, authored by scientists and evaluated by journal editors, program chairs, and peer reviewers, edited by journal staff
- *Publications*, authored by scientists, read and assessed by other scientists, systematic reviewers, database curators, funding agencies, IRBs, academic institutions, and policymakers

Clark *et al.* (2014) conceptualize the ecosystem of biomedical communication as a cycle of nine activities, with inputs and outputs (the output of the last activity feeding back into the first):

1. Authoring
2. Reviewing for Publication
3. Editing and Publishing
4. Database and Knowledge-base Curation
5. Searching for Information
6. Reading
7. Discussion
8. Evaluating and Integrating Information
9. Experiment Design and Execution

Textual artifacts are primary inputs and outputs of some of these activities. For example, inputs for authoring include other relevant publications in the research space, as well as experimental data and observations, and the output is a manuscript for submission. By providing the ability to automatically process such artifacts at a large scale and extract relevant information for subsequent activities, bioNLP methods have the potential to assist scientists with the entire lifecycle of biomedical communication. Though not explicit in Clark *et al.*’s conceptualization, the same capabilities can also benefit other stakeholders (journal editors, reviewers, funding agencies, etc.), who need to evaluate such artifacts based on their scientific merit.

What kinds of text mining tools can be envisioned? What kinds of benefits can they provide? We briefly outline several categories of tools and their potential benefits below.

1. *Plagiarism/fraud detection:* Although plagiarism and outright fraud are relatively rare (though seemingly growing (Fang *et al.*, 2012)) in scientific literature, tools that can detect plagiarism/fraud can be helpful to journal editors in catching research misconduct before publishing an article and avoiding later retractions, which may reflect badly on the journal.
2. *Adherence to reporting guidelines:* Tools that can assess a manuscript against the relevant reporting guidelines and flag the issues would be useful for journal editors, who can then require the authors to fix the problems for publication.
3. *Managing information overload:* Text mining tools can help in managing information overload by summarizing and aggregating salient knowledge (e.g., hypotheses, claims, supporting evidence) in textual artifacts, a capability that can benefit all stakeholders. Efficient knowledge management can help research rigor and reduce research waste by ensuring that, for example, scientists are aware of all relevant studies before embarking on a research project (Lund *et al.*, 2016) or that funding agencies are not awarding funds to redundant or unjustified proposals (Robinson and Goodman, 2011; Habre *et al.*, 2014).
4. *Accurate citation and enhanced bibliometrics:* Tools that can verify whether the authors cite relevant literature (or omit) accurately would be beneficial in reducing citation distortion, which has been shown to lead to unfounded authority (Greenberg, 2009). Advanced citation analysis tools that can recognize the function of a citation and its significance for the publication can help funding agencies and academic institutions move beyond simple citation counts and make more informed decisions about the impact of a particular study.

Although text mining has been used to address a variety of tasks that can be subsumed by the categories outlined above, there is little research on using text mining for broadly addressing research integrity and rigor issues in biomedical science. One nascent effort is a collaboration between the academic publisher Elsevier and Humboldt University^3^, which aims to use text/data mining for early detection of integrity issues, focusing mainly on plagiarism/fraud, image manipulation, and data fabrication, although no published results were available at the time of this writing.

In the remainder of this section, we review the existing NLP/bioNLP research on the four categories of tasks that we outlined above.

### 2.1 Plagiarism/Fraud Detection

Plagiarism is “the appropriation of another person’s ideas, processes, results, or words without giving appropriate credit” (Habibzadeh and Shashok, 2011). A serious problem in academia, especially with regards to student writing, plagiarism also occurs in medical publications (Habibzadeh and Shashok, 2011). Plagiarism comes in several forms: at one end of the spectrum is copy-paste plagiarism, which is relatively easy to detect, and on the other end is paraphrased or translated plagiarism, which can be very challenging. Plagiarism detection is a well-studied information retrieval task and dedicated tools have been developed (e.g., TurnItIn^4^, Plagramme^5^). CrossRef Similarity Check^6^, a TurnItIn-based tool used by publishers, specifically targets plagiarism in scholarly communication. It generates an overall similarity score between a manuscript and the articles in a large database of publications and flags the manuscript if its similarity score is over a publisher-determined threshold.

Generally, two plagiarism detection tasks are distinguished: *extrinsic* and *intrinsic plagiarism detection*. In *extrinsic plagiarism detection*, a document is compared to other candidate documents in a reference collection. Approaches to this task differ with respect to how documents are represented: document fingerprints based on substring hashing (Hoad and Zobel, 2003) or vectors (Stein and Meyer zu Eissen, 2006). Vector representations can be based on characters, words, word sequences (n-grams), or stopwords (Stamatatos, 2011). High number of candidate documents can pose challenges in extrinsic plagiarism detection (Stein and Meyer zu Eissen, 2006); therefore, efficient document representations and candidate document selection can be critical. In *intrinsic plagiarism detection*, the goal is to recognize shifting writing styles within a document to spot plagiarism (Meyer zu Eissen and Stein, 2006). Methods for this task rely on stylometric features, such as word class usage and average word frequency, which indicate the author’s vocabulary size and style complexity. Plagiarism detection has been the topic of PAN shared task challenges^7^, the last edition of which took place in 2016 (Rosso *et al.*, 2016). Performance for extrinsic plagiarism detection in these competitions has reached an F_1_ score of 0.88 (Sanchez-Perez *et al.*, 2014), while the state-of-the-art performance for intrinsic plagiarism detection is much lower at an F_1_ score of 0.22 (Kuznetsov *et al.*, 2016). Plagiarism in the PAN corpora was simulated, whereas Nawab *et al.* (2016) used a corpus of PubMed abstracts that were deemed to be suspiciously similar to other abstracts and used a query expansion approach based on UMLS Metathesaurus (Lindberg *et al.*, 1993) to detect plagiarism. Plagiarism detection tools are most beneficial to journal editors and peer reviewers, though scientists can also benefit from using such tools to prevent inadvertent plagiarism or self-plagiarism.

Plagiarism accounts for a relatively small fraction of retractions in biomedical research articles (9.8%), while the fraud accounts for 43.4% (Fang *et al.*, 2012). Such cases often involve data fabrication or falsification (Fanelli, 2009), types of misconduct that would typically be difficult to flag with text analysis alone. Focusing on text only, Markowitz and Hancock (2015) investigated whether scientists write differently when reporting fraudulent research. They compared the linguistic style of publications retracted for fraudulent data with that of unretracted articles and articles retracted for reasons other than fraud. They calculated a linguistic obfuscation score based on stylistic and psycholinguistic characteristics of a document, including ease of reading, rate of jargon, causal and abstract words. They found that retracted articles were written with significantly higher levels of linguistic obfuscation and that obfuscation was positively correlated with the number of references. However, their score-based method had a high false-positive rate and they suggested that NLP techniques could achieve higher classification performance. A task similar to fraud detection, considered in open-domain NLP, is *deception detection*, generally cast as a binary classification task (deceptive vs. true) (Mihalcea and Strapparava, 2009). Supervised machine learning techniques (Support Vector Machines (SVMs), Naive Bayes) using n-gram and psycholinguistic features have been applied to this task (Mihalcea and Strapparava, 2009; Ott *et al.*, 2011), the latter achieving F_1_ score of 0.90 (Ott *et al.*, 2011). Interestingly, inter-annotator agreement (Fleiss’ κ=0.11) and human judgements (0.60 F_1_) were found to be lower than machine performance.

The classification approach used for deception detection is likely to be beneficial in detecting fraudulent articles. Similarly to plagiarism detection, fraud detection tools would be most useful to journal editors. We note that, in general, decisions regarding fraud or plagiarism should ultimately only be made by humans, since such accusations can be damaging to a scientist’s career. However, text mining approaches can, to some extent, flag suspicious manuscripts, which can then be given extra scrutiny.

### 2.2 Adherence to Reporting Standards and Guidelines

The EQUATOR Network (Simera *et al.*, 2010) has promoted transparent and accurate reporting, and guidelines for various study types have been developed under its umbrella (e.g., CONSORT for randomized clinical trials (RCTs) (Schulz *et al.*, 2010), ARRIVE for preclinical animal studies (Kilkenny *et al.*, 2010)). For example, the CONSORT Statement consists of a 25-item checklist and a flow diagram. The CONSORT checklist for Methods sections is provided in the Supplementary File as an example.

Although a large number of journals have adopted or support such guidelines, adherence to them remains inadequate (Turner *et al.*, 2012). As a solution, structured reporting of key methods and findings has been proposed (Altman, 2015). Until such proposals gain currency, however, most methodological information is likely to remain buried in narrative text or, in the worst case, be completely absent from the publication. Text mining tools can help journal editors enforce adherence to reporting guidelines by locating key statements corresponding to specific guideline items or giving alerts in their absence. For example, per CONSORT, the method used to generate the random allocation sequence as well as statistical details critical for reproducibility can be identified. Recognition of description of limitations and sources of possible bias can be beneficial for broader rigor and generalizability. Additionally, medical journals require or encourage inclusion of certain types of meta-information, such as funding, conflicts of interest, trial registration, and data access statements. Identifying such meta-statements and locating statements corresponding to guideline items can both be categorized as *information extraction* tasks, and similar techniques (text/sentence classification, sequence labeling) can be applied to them. The difficulty of extracting these items varies widely: locating trial registration information seems relatively easy, since the trial registration numbers have a standard format, whereas extracting statements that indicate that interpretation is “consistent with results, balancing benefits and harms, and considering other relevant evidence” (a CONSORT checklist item) seems challenging, since some subjectivity may be involved and a deeper text understanding may be needed. Commercial software has been developed to address adherence issues to some extent (e.g., Penelope Research^8^, StatReviewer^9^); however, they currently have limited capabilities and information about the underlying technologies is sparse. Tools that can address the guideline adherence would be useful not only for journals and reviewers, but also to authors of systematic reviews, who aim to identify well-designed, rigorous studies, and to clinicians looking for the best available clinical evidence.

We are not aware of any published bioNLP research that aims to determine whether a manuscript fully complies with the relevant set of reporting guidelines. However, some research attempts to identify some guideline items as well as other meta-information, often for the purpose of automating systematic review process (Tsafnat *et al.*, 2014). We discuss such research below; see O’Mara-Eves *et al.* (2015) for a general discussion of using text mining for systematic reviews.

In the simplest case, some statistical details, such as *p*-values and confidence intervals, can be identified with simple regular expressions (Chavalarias *et al.*, 2016). Some meta-information, such as funding, conflict of interest, or trial registration statements, often appear in dedicated sections and are expressed using a limited vocabulary; hence, simple keyword-based techniques could be sufficient. For example, Kafkas *et al.* (2013) mined data accession numbers in full-text articles using regular expressions.

Other items require more sophisticated techniques, often involving machine learning. For example, Kiritchenko *et al.* (2010) extracted 21 key elements (e.g., eligibility criteria, sample size, primary outcomes, registration number) from full-text RCT articles, some of which are included in CONSORT. Their system, trained on an annotated corpus of 132 articles, used a two-stage pipeline: first, given a particular element, a classifier predicted whether a sentence is likely to contain information about that element. Second, regular expressions were used to extract the exact mention of the element. Some meta-information (DOI, author name, publication date) was simply extracted from PubMed records. For each remaining element, an SVM model with n-gram features was trained for sentence classification. A post-hoc evaluation on a test corpus of 50 articles yielded a precision of 0.66 for these elements (0.94 when partial matches were considered correct).

Névéol *et al.* (2011a) focused on extracting deposition statements of biological data (e.g., gene sequences) from full-text articles. They semi-automatically constructed a gold standard corpus. Their approach consisted of two machine learning models: one recognized data deposition components (Data, Action, General Location, Specific Location) using the Conditional Random Fields (CRF) sequence labeling algorithm. The main model, a binary classifier, predicted whether a sentence contains a data deposition statement. This classifier, trained with Naive Bayes and SVM algorithms, employed token, part-of-speech, and positional features as well as whether the sentence included components identified by the CRF model. An article was considered positive for data deposition if the top-scored sentence was classified as positive. Their system yielded an F_1_ score of 0.81.

PICO frame elements (*Problem*, *Population*, *Intervention*, *Comparison*, and *Outcome*) are recommended for formulating clinical queries in evidence-based medicine (Sackett *et al.*, 1996). They often appear in reporting guideline checklists (e.g., Participants in CONSORT vs. *Population* in PICO). Some research focused on PICO and its variants. Demner-Fushman and Lin (2007) identified PICO elements in PubMed abstracts for clinical question answering. Outcomes were extracted as full sentences and other elements as short noun phrases. The results from an ensemble of classifiers (rule-based, n-gram-based, position-based, and semantic group-based), trained on an annotated corpus of 275 abstracts, were combined to recognize outcomes. Other elements were extracted using rules based on the output of MetaMap (Aronson and Lang, 2010), a system that maps free text to UMLS Metathesaurus concepts (Lindberg *et al.*, 1993). Recently, Wallace *et al.* (2016), noting that PICO elements may not appear in abstracts, attempted to extract PICO sentences from full-text RCT articles. They generated sentence-level annotations automatically from free-text summaries of PICO elements in the Cochrane Database of Systematic Reviews (CDSR), using a novel technique called *supervised distant supervision*. A small number of sentences in the original articles that were most similar to CDSR summary sentences were identified and a manually annotated subset was leveraged to align unlabeled instances with the structured data in CDSR. Separate models were learned for each PICO element with bag-of-words and positional features as well as features regarding the fraction of numerical tokens, whether the sentence contains a drug name, among others. Their technique outperformed models that used direct supervision or distant supervision only. A PICO variant, called PIBOSO (B: Background, S: Study design, O: Other) was also studied (Kim *et al.*, 2011). A corpus of 1,000 PubMed abstracts were annotated with these elements (PIBOSO-NICTA) and two classifiers were trained on this corpus: one identified PIBOSO sentences and the other assigned PIBOSO labels to these sentences. A CRF model was trained using bag-of-words, n-gram, part-of-speech, section heading, position, and sequence features as well as domain information from MetaMap. With the availability of the PIBOSO-NICTA corpus, several studies explored similar machine-learning based approaches (Verbeke *et al.*, 2012; Hassanzadeh *et al.*, 2014), state-of-the-art results, without using any external knowledge, were reported by Hassanzadeh *et al.* (2014) (0.91 and 0.87 F_1_ scores on structured and unstructured abstracts, respectively).

Marshall *et al.* (2015) developed a tool called RobotReviewer to identify risk of bias (RoB) statements in clinical trials. They used seven risk categories specified in the Cochrane RoB tool (e.g., random sequence generation, allocation concealment, blinding of participants and personnel, and selective outcome reporting) and labeled articles as high or low risk with respect to a particular category. Similar to their approach for extracting PICO statements, they semi-automatically generated positive instances for training by leveraging CDSR, where systematic reviewers copy/paste a fragment from the article text to support their RoB judgements. An SVM classifier based on multi-task learning mapped articles to RoB assessments and simultaneously extracted supporting sentences with an accuracy of 0.71 (compared to 0.78 human accuracy).

Considering research rigor broadly, Kilicoglu *et al.* (2009) developed machine learning models to recognize methodologically rigorous, clinically relevant publications to serve evidence-based medicine. Several binary classifiers (Naive Bayes, SVM, and boosting) as well as ensemble methods (stacking) were trained on a large set of PubMed abstracts previously annotated to develop PubMed Clinical Queries filter (Wilczynski *et al.*, 2005). The base features used included token, PubMed metadata as well as semantic features, as extracted by MetaMap and SemRep (Rindflesch and Fiszman, 2003), a biomedical relation extraction tool. Best results (F_1_ score of 0.67) were achieved with a stacking classifier that used base models trained with various feature-classifier combinations (e.g., SVM with token features only).

### 2.3 Managing Information Overload

Tasks we discussed so far did not require much deep natural language understanding; surface-level features, such as n-grams and part-of-speech tags, positional information and limited semantic knowledge, extracted with tools like MetaMap, were mostly sufficient for models that yielded reasonable performance. In this section, we turn to tasks aiming to address information overload caused by the considerable size and the rapid growth of the biomedical literature.

A strategy for efficient management of the biomedical literature should support extraction of the hypotheses and the key arguments made in a research article (referred to as *knowledge claims* (Myers, 1992) henceforth) as well as their contextualization (e.g., identifying the evidence provided to support these claims, the level of certainty with which the claims are expressed, and whether they are new knowledge). It should also allow aggregating such knowledge over the entire biomedical literature. A deeper text understanding is required for such capabilities and we argue that the key to them is *normalization* of claims and the supporting evidence into computable semantic representations that can account for lexical variability and ambiguity. Such representations make the knowledge expressed in natural language amenable to automated inference and reasoning (Blackburn and Bos, 2005). Furthermore, they can form the building blocks for advanced information seeking and knowledge management tools, such as semantic search engines, which can help us navigate the relevant literature more efficiently. For example, formal representations of knowledge claims can underpin tools that enable searching, verifying, and tracking claims at a large scale, and summarizing research on a given biomedical topic; thus, reducing the time spent locating/retrieving information and increasing the time spent interpreting it. Such tools can also address siloization of research (Swanson, 1986; Editorial, 2016), putting research questions in a larger biomedical context and potentially uncovering previously unknown links from areas that the researcher does not typically interact with. Literature-scale knowledge extraction and aggregation on a continuous basis can also facilitate ongoing literature surveillance, with tools that alert the user when a new knowledge claim related to a topic of interest is made, when a claim of interest to the user is discredited or contradicted^10^, increasing research efficiency. Advanced knowledge management tools would be beneficial to all parties involved in biomedical research: i) to researchers in keeping abreast of the literature, generating novel hypotheses, and authoring papers, ii) to funding agencies, IRBs, and policymakers in better understanding the state-of-the-art in specific research areas, creating research agendas/policies, verifying claims and evidence presented in proposals, assessing whether the proposed research is justified, iii) to journal editors, peer reviewers, systematic reviewers, and database curators in locating, verifying, and tracking claims and judging evidence presented in manuscripts and publications.

What do we mean by normalization of knowledge claims and evidence? With normalization, we refer to recognition of biomedical entities, their properties, and the relationships between them expressed in text and mapping them to entries in a relevant ontology or knowledge-base. As the basis of such formalization, we distinguish three levels of semantic information to be extracted: *conceptual*, *relational*, and *contextual*. Roughly, the conceptual level is concerned with biomedical entities (e.g., diseases, drugs), relational level with biomedical relationships (e.g., gene-disease associations), and the contextual level with how these relationships are contextualized and related for argumentation. A knowledge claim, in the simplest form, can be viewed as a relation. We illustrate these levels on a PubMed abstract in the Supplementary File.

Conceptual level is in the purview of the *named entity recognition and normalization* (NER/NEN) task, while *relation extraction* focuses on the relational level. These tasks are well-studied in bioNLP. We provide a brief overview in the Supplementary File; see recent surveys (Gonzalez *et al.*, 2016; Luo *et al.*, 2016) for more comprehensive discussion. In the remainder of this subsection, we first briefly discuss tools that address information overload using concepts and relations extracted from the literature and then turn to research focusing on the contextual level.

#### 2.3.1 Literature-scale relation extraction

Literature-scale relation extraction has been proposed as a method for managing information overload (Kilicoglu *et al.*, 2008). SemMedDB (Kilicoglu *et al.*, 2012) is a database of semantic relations extracted with SemRep (Rindflesch and Fiszman, 2003) from the entire PubMed. In its latest release (as of December 31th, 2016), it contains about 89 million relations extracted from more than 26 million abstracts. It has been used for a variety of tasks, such as clinical decision support (Jonnalagadda *et al.*, 2013), uncovering potential drug interactions in clinical data (Zhang *et al.*, 2014), supporting gene regulatory network construction (Chen *et al.*, 2014), and medical question answering (Hristovski *et al.*, 2015). It also forms the back-end for the Semantic MEDLINE application (Kilicoglu *et al.*, 2008), which integrates semantic relations with automatic abstractive summarization (Fiszman *et al.*, 2004), and visualization, to enable the user navigate biomedical literature through concepts and their relations. Semantic MEDLINE, coupled with a literature-based discovery extension called “discovery browsing”, was used to propose a mechanistic link between age-related hormonal changes and sleep quality (Miller *et al.*, 2012) and to elucidate the paradox that obesity is beneficial in critical care despite contributing to disease generally (“the obesity paradox”) (Cairelli *et al.*, 2013). Another database, EVEX (Van Landeghem *et al.*, 2013), is based on the Turku Event Extraction System (TEES) (Björne and Salakoski, 2011) and includes relations extracted from the full-text articles in PMCOA as well as PubMed abstracts. It consists of approximately 40 million bio-molecular events (e.g., gene expression, binding). A CytoScape plugin, called CyEVEX, is made available for integration of literature analysis with network analysis. EVEX has been exploited for gene regulatory network construction (Hakala *et al.*, 2015). Other databases, such as PharmGKB (Hewett *et al.*, 2002) and DisGeNET (Piñero *et al.*, 2015), integrate relationships extracted with text mining with those from curated resources.

#### 2.3.2 Contextualizing Biomedical Relations

Contextualizing relations (or claims) focuses on how they are presented and how they behave in the larger discourse. Two distinct approaches can be distinguished.

The first approach, which can be considered “bottom-up”, focuses on classifying scientific statements or relations along one or more *meta-dimensions* aiming to capture their contextual properties; for example, whether they are expressed as speculation or not. One early task adopting this approach was distinguishing speculative statements from facts (*hedge classification*). For this task, weakly supervised learning techniques (Szarvas, 2008), as well rule-based methods using lexical and syntactic templates (Kilicoglu and Bergler, 2008; Malhotra *et al.*, 2013) have been explored, yielding similar performance (0.85 F_1_ score)^11^. Semantically more fine-grained, speculation/negation detection task has focused on recognizing speculation and negation cues in text (e.g., *suggest*, *likely*, *failure*) and their linguistic scope, often formalized as a relation (Kim *et al.*, 2008) or a text segment (Vincze *et al.*, 2008). Speculation/negation detection has been studied in the context of BioNLP Shared Tasks on event extraction (Kim *et al.*, 2009, 2012) and the CoNLL’10 Shared Task on Hedge Detection (Farkas *et al.*, 2010). Supervised machine learning methods (Björne and Salakoski, 2011; Morante *et al.*, 2010) as well as rule-based methods with lexico-syntactic patterns (Kilicoglu and Bergler, 2012) have been applied to this task. The interaction of speculation and negation has been studied under the notion of *factuality* and factuality values (`Fact, Probable, Possible, Doubtful, Counterfact`) of biological events were computed using a rule-based, syntactic composition approach (Kilicoglu *et al.*, 2015).

Focusing on a more comprehensive characterization of scientific statements, Wilbur *et al.* (2006) categorized sentence segments along five dimensions: Focus (whether the segment describes a finding, a method, or general knowledge), Polarity (positive/negative), Certainty (the degree of speculativeness expressed towards the segment on a scale of 0-3), Evidence (four levels, from no stated evidence to explicit experimental evidence in text), and Direction (whether segment describes an increase or decrease in the finding). A similar categorization (“meta-knowledge”) was proposed by Thompson *et al.* (2011), who applied it to events, rather than arbitrary text segments. They also proposed two hyper-dimensions that are inferred from their five categories: one indicates whether the event in question is New Knowledge and the other whether it is a Hypothesis. Studies that focused on predicting these meta-dimensions have been trained on the annotated corpora and used supervised machine learning techniques (Shatkay *et al.*, 2008; Miwa *et al.*, 2012b). The Claim Framework (Blake, 2009) proposed a categorization of scientific claims according to the specificity of evidence, somewhat similar to Focus dimension in the schema of Wilbur *et al.* (2006). Five categories were distinguished (explicit claim, implicit claim, observation, correlation, and comparison). A small corpus of full-text articles was annotated with these categories and an approach based on lexico-syntactic patterns was used to recognize explicit claims.

The second approach (“top-down”) focuses on classifying larger units (sentences or a sequence of sentences) according to the function they serve in the larger argumentative structure. Proposed by Teufel *et al.* (1999, 2009) for scientific literature on computational linguistics and chemistry, *argumentative zoning* assigns sentences to domain-independent zone categories based on the rhetorical moves of global argumentation and the connections between the current work and the cited research. The proposed categories include, for example, `Aim` (statement of specific research goal or hypothesis), `Nov Adv` (novelty/advantage of the approach), `Own_Mthd` (description of methods used), among others. Mizuta *et al.* (2006) adapted this classification to biology articles and presented an annotated corpus. Guo *et al.* (2011) adopted a simplified version of argumentative zoning with seven classes (e.g., Background, Method, Result, and Future Work). They used weakly supervised SVM and CRF models to classify sentences in abstracts discussing cancer risk assessment, which yielded an accuracy of 0.81. The CoreSC schema (Liakata *et al.*, 2010) is an extension of the argumentative zoning approach, in which sentences are classified along two layers according to their role in scientific investigation. The first layers consists of 11 categories (e.g., Background, Motivation, Experiment, Model, Result, Conclusion) and the second layer indicates whether the information is New or Old. A corpus of chemistry articles annotated with these layers was presented. SVM and CRF classifiers that recognize the first layer categories were developed (Liakata *et al.*, 2012a), achieving best results with Experiment, Background, and Model classes (0.76, 0.62, 0.53 F_1_ scores, respectively). N-gram, dependency, and document structure features (section headings) were found to be predictive. Such top-down classifications are similar to but more fine-grained than IMRaD rhetorical categories (Introduction, Methods, Results, Discussion) that underlie the structure of most scientific articles. Since the sentences may not conform to the characteristics of the section that they appear in, some research considered classifying sentences into IMRaD categories. For example, Agarwal and Yu (2009) compared several rule-based and supervised learning methods to classify sentences from full-text biomedical articles into these categories. The best results reported (0.92 accuracy and F_1_ score) were obtained with a Naive Bayes classifier with n-gram, tense, and citation features, and feature selection. Other similar categorizations have also been proposed (e.g., (de Waard *et al.*, 2009)). Note that the methods applied in these approaches are largely similar to those discussed earlier for identification of specific statements, such as PICO or data deposition statements. Finally, a comprehensive, multi-level model of scientific argumentation, called Knowledge Claim Discourse Model (KCDM), has been proposed by Teufel (2010). Five levels varying in their degree of abstraction have been distinguished. At the most abstract level, *rhetorical goals* are formalized into four categories, often not explicit in text (“Knowledge claim is significant”, “Knowledge claim is novel”, “Authors are knowledgeable”, “Research is methodologically sound”). Next level, *rhetorical moves*, addresses the properties of the research space (e.g.,”No solution to new problem exists”) and the new knowledge claim (e.g., “New solution solves problem”). The third level, *knowledge claim attribution*, is concerned with whether a knowledge claim is attributed to the author or others. At the fourth level are *hinge moves*, which categorize the connections between the new knowledge claim and other claims (e.g., “New claim contrasts with existing claim”). The bottom and the most concrete layer, *linearization and presentation*, deals with how these rhetorical elements are realized within the structure of the article. Teufel reported the results of several annotation studies focusing on argumentative zoning and knowledge claim attribution (κ values of 0.71 to 0.78), and her argumentative zone detection system, based on supervised learning with verb features, word lists, positional information, and attribution features, achieved a *κ* value 0.48, with respect to the annotated corpus.

Similarly taking a top-down approach but focusing on the relations between individual discourse segments (similar to KCDM *hinge moves*) are models of *discourse coherence*. Such relations include elaboration, comparison, contrast, and precedence and are often indicated with discourse connectives (e.g., *furthermore*, *in contrast*). Linguistic theories and treebanks have been proposed to address these relations, including Rhetorical Structure Theory (RST) (Mann and Thompson, 1988) and the Penn Discourse TreeBank (PDTB) (Miltsakaki *et al.*, 2004), each assuming a somewhat different discourse structure and relation inventory and differing in their level of formalization. In the biomedical domain, discourse relations remain understudied, with the notable exception of the Biomedical Discourse Relation Bank (BioDRB) corpus (Prasad *et al.*, 2011), in which a subset of PDTB relation types were used to annotate abstracts in the GENIA corpus. Detection of discourse connectives was explored on this corpus and an F_1_ score of 0.76 was achieved with a supervised learning approach and domain adaptation techniques (Ramesh *et al.*, 2012).

Some research considered combining bottom-up and top-down approaches for a fuller understanding of scientific discourse or contextual meaning. For example, a three-way characterization, based on meta-knowledge dimensions (Thompson *et al.*, 2011), CoreSC (Liakata *et al.*, 2010), and discourse segment classification (de Waard *et al.*, 2009), was attempted and these components were shown to be complementary (Liakata *et al.*, 2012b). The Embedding Framework (Kilicoglu, 2012) proposed a unified, domain-independent semantic model for contextual meaning, consolidating the meta-dimensions and discourse coherence relations. A fine-grained categorization of contextual predicates was presented, with 4 top-level categories (`Modal, Valence Shifter, Relational, Propositional`), where the `Modal` and `Valence_Shifter` categories overlap with meta-dimensions and the Relational category overlaps with discourse relations. A dictionary of terms, classified according to this fine-grained categorization, was constructed and a rule-based interpretation methodology based on this dictionary and syntactic dependency composition was proposed. The framework is designed to complement existing relation extraction systems. While no specific corpus annotation was performed, the methodology has been applied to relevant tasks, such as speculation/negation detection (Kilicoglu and Bergler, 2012), factuality assessment (Kilicoglu *et al.*, 2015), and attribution detection (Kilicoglu, 2012), yielding good performance.

Although not a text mining approach, an effort that deserves discussion here is Micropublications (Clark *et al.*, 2014), a semantic model of scientific claims, evidence, and arguments. Built on top of Semantic Web technologies, micropublications are intended for use in the research lifecycle, where scientists create, publish, expand, and comment on micropublications for scientific communication. They have been proposed as a potential solution to improve research reproducibility and robustness. At a minimum, a micropublication is conceived as a claim with its attribution, and in its full form, as a claim with a complete directed-acyclic support graph, consisting of relevant evidence, interpretations, and discussion that supports/refutes the claim, either within the publication or in a network of publications discussing the claim. It has been designed to be compatible with claim-based models formalizing relationships (e.g., Nanopublications (Mons and Velterop, 2009)), as well as with claims in natural language text. The model can accommodate artifacts such as figures, tables, images, and datasets, which text mining approaches generally do not consider. While it has been used for manual annotation (Schneider *et al.*, 2014), to our knowledge, the Micropublications model has not been used as a target for text mining. An example micropublication is presented in the Supplementary File.

### 2.4 Accurate Citation and Enhanced Bibliometrics

Citations are important for several reasons in ensuring research integrity/rigor. First, the performance of a scientist is often measured by the number of citations they receive and the number of articles they publish in high impact-factor journals. Count-based measures, such as the h-index (Hirsch, 2005), are often criticized, because they treat all citations as equal and do not distinguish between the various ways and reasons a paper can be cited. For example, a paper can be appraised in a positive light or criticized, it can be cited as the basis of the current study or more peripherally. Such differences should be accounted for enhanced bibliometric measures. More sophisticated measures have been proposed in response to such criticism (e.g., (Hutchins *et al.*, 2016)). Secondly, from an integrity perspective, it is important to ensure that all references in a manuscript (or any other scientific textual artifact) are accurately cited. Two kinds of reference accuracy problems are distinguished (Wager and Middleton, 2008): *citation accuracy* refers to the accuracy of details, such as authors’ names, date of publication, and volume number, whereas *quotation accuracy* refers to whether the statements from the cited papers are accurately reflected in the citing paper. Reference accuracy studies were found to have a median error rate of 39% and quotation accuracy studies a median error rate of 20% (Wager and Middleton, 2008). Greenberg (2009) highlighted some types of citation distortions (i.e., quotation accuracy problems) that lead to unfounded authority. For example, *citation transmutation* refers to “the conversion of hypothesis into fact through act of citation alone”, and *dead-end citation* to “citation to papers that do not contain content addressing the claim.” Another rigor issue is the continued citation of retracted papers, which may lead to spreading of misinformation. A study of retracted paper citations found that 94% of the citing papers did not acknowledge the retraction (Budd *et al.*, 2011). Automated citation analysis tools and accuracy checkers would be beneficial for journal editors and staff in their workflows, as well as for scientists in authoring manuscripts and academic institutions and funding agencies in considering quality of impact rather than quantity and improving decision-making.

Most text mining research on citations has focused on the computational linguistics literature, an area in which a corpus of full-text articles is available (ACL Anthology Corpus (Radev *et al.*, 2009)). Citation analysis has been proposed for enhancing bibliometrics as well as for extractive summarization (Qazvinian and Radev, 2008). Several aspects of citations have been studied. Research on *citation context detection* aims to identify the precise span of the discussion of the reference paper in the citing paper. For example, to detect the surrounding sentences that discuss a reference paper, Qazvinian and Radev (2010) proposed a method based on Markov Random Fields using sentence similarity and lexical features from sentences. Abu-Jbara and Radev (2012) focused on reference scopes that are shorter than the full sentence. They explored several methods for this task: word classification with SVM and logistic regression, CRF-based sequence labeling, and segment classification which uses rules based on CRF results, achieving best performance with segment classification (F_1_ score of 0.87). Other studies explored *citation significance*. Athar (2014) presented a text classification approach to determine whether a citation is significant for the citing paper and achieved 0.55 F_1_ score with a Naive Bayes classifier that used as features number of sentences with acronyms, with formal citation to the paper and to the author’s name, as well as average similarity of the sentence with the title. Similar text classification techniques were used to identify key references (Zhu *et al.*, 2015) and meaningful citations (Valenzuela *et al.*, 2015); the number of times a paper is cited was identified as the most predictive feature. *Citation sentiment* (whether the authors cite a paper positively, negatively, or neutrally) has also been proposed to enhance bibliometrics. Athar (2011) annotated the ACL Anthology Corpus for citation sentiment and used an SVM classifier with n-gram and dependency features extracted from the citation sentence for sentiment classification, achieving a macro-F_1_ score of 0.76. In the biomedical domain, Xu *et al.* (2015) annotated the discussion sections of 285 RCT articles with citation sentiment. Using an SVM classifier with n-gram and various lexicon-based features (e.g., lexicons of positive/negative sentiment, contrast expressions), they reached a macro-F_1_ score of 0.72. A more fine-grained citation classification concerns *citation function*, for which many classifications have been proposed. For example, Teufel *et al.* (2006a) presented a scheme, which contained 12 categories (e.g., `Weak` (weakness of the cited approach), `PBas` (cited work as the starting point), `CoCo-` (unfavorable comparison/contrast)) and measured inter-annotator agreement (κ=0.72). Later, Teufel *et al.* (2006b) used a memory-based learning algorithm to recognize these categories. They used features based on cue phrases in the citation sentence, position of the citation, and self-citation, which yielded a κ of 0.57. In the biomedical domain, Agarwal *et al.* (2010) presented a corpus of 43 biomedical articles annotated with eight citation roles (e.g., Background/Perfunctory, Contemporary, Contrast/Conflict, Evaluation, Modality, Similarity/Consistency), achieving moderate inter-annotator agreement (κ=0.63), though it seems difficult to think of some of their categories (Contemporary, Modality) as citation roles in a traditional sense. Using n-gram features with SVM and Naive Bayes classifiers, they obtained a macro-F_1_ score of 0.75.

The first type of reference accuracy, referred to as *citation accuracy* above, is studied under the rubric of *citation matching*. We do not discuss this task here, as NLP has little relevance to it; see Olensky *et al.* (2016) for a comparison of several citation matching algorithms. Ensuring quotation accuracy, on the other hand, can be viewed as a text mining task, in which the goal is to identify the segments of the reference paper that are discussed in the citing paper. Inability to find such a segment would indicate a *dead-end citation*, while finding inconsistencies between how a claim is presented in the reference paper versus the citing paper with respect to its factuality might indicate a *citation transmutation* (Greenberg, 2009). However, identifying reference paper segments precisely can be challenging, as the citing paper usually does not simply quote the reference paper verbatim, but rather paraphrases its contents, and commonly, refers to its contents in an abstract manner. In the Text Analysis Conference (TAC) 2014 Biomedical Summarization shared task^12^, one subtask involved finding the spans of text in reference papers that most accurately reflect the citation sentence and identifying what facet of the reference paper it belongs to (e.g., Hypothesis, Method, Results, Implication). The task focused on a corpus of 20 biology articles, each with 10 corresponding reference articles. The inter-annotator agreement was found to be low. The results of this shared task were not available at the time of this writing; however, one of the reported systems (Molla *et al.*, 2014) relied on calculating text similarity between the citation sentence and the sentences in the reference paper, using *tf.idf*, as well as various methods to expand the citation context and the reference paper context for similarity calculation. The best results (F_1_ score of 0.32) were obtained when using 50 sentences surrounding the citation sentence and all sentences from the articles that cite the reference paper for context. The same task has also been adapted to the computational linguistics literature (Jaidka *et al.*, 2016); even though the results have been poorer, with the top-ranking system obtaining 0.1 F_1_ score (Cao *et al.*, 2016).

## 3 Challenges and Directions

We examined four areas of concern for biomedical research integrity and rigor and discussed existing text mining research that has the potential to address them. We discuss below several general challenges facing bioNLP research focusing on these areas and highlight some promising avenues for future research.

The first challenge is concerned with availability of artifacts that can be used to train text mining methods. While most text mining research has focused on PubMed abstracts due to their availability, most biomedical knowledge relevant to the tasks discussed, including study details, knowledge claims, and citations, can only be located in full-text. Blake (2009) found that only 8% of the explicit claims were expressed in abstracts. Furthermore, biomedical abstracts differ from full-text in terms of structure and content (Cohen *et al.*, 2010). The PMC-OA subset is amenable to automated approaches without much additional pre-processing effort; however, it contains only about a million full-text articles (4% of all PubMed abstracts). Due to availability and access difficulties, researchers often use non-standard PDF-to-text conversion tools to extract full-text from PDF files (e.g., (Marshall *et al.*, 2015; Jimeno-Yepes and Verspoor, 2014)). Considering that the progress in bioNLP is partly attributed to public accessibility of biomedical abstracts, a similar mode of access can further stimulate research in mining of full-text articles. We are not aware of research focusing on other textual artifacts discussed, though abstracts of NIH grant applications and the resulting publications are available via NIH RePORT^13^ and some journals (e.g., British Medical Journal) publish pre-publication manuscripts and reviewer reports for transparency.

Collecting bibliographic data at a large scale also remains challenging. Two sources of scholarly citation considered most authoritative, Web of Science and Scopus, are neither complete nor fully accurate (Franceschini *et al.*, 2016) and require high subscription fees. Others, like Google Scholar, have license restrictions. Open Citations Corpus (OCC) has been proposed as an open-access repository of citation data to improve citation access (Peroni *et al.*, 2015). They rely on the SPAR ontologies (Peroni, 2014), which define characteristics of the publishing domain. Citation information in PMC-OA has been made available in OCC. Although this is a small subset of the biomedical literature, the movement towards open-access citation data is encouraging for research.

Even when the text sources are plentiful, restrictions may apply to text mining of their contents. Publishers often adopt a license-based approach, allowing researchers from subscribing institutions to register for an API key to text-mine for research purposes. Negotiating a separate license with each publisher is not only impractical for both researchers and publishers but also ineffective, since some tasks (e.g., plagiarism detection, managing information overload, citation analysis) presuppose text mining at the literature scale with no publisher restrictions. The Crossref Metadata API initiative (Lammey, 2016) aims to solve this problem by providing direct links to full-text on the publisher’s site and a common mechanism for recording license information in Crossref metadata. Several publishers (e.g., HighWire Press, Elsevier, Wiley) as well as professional societies (e.g., American Psychological Association) have been involved in this initiative.

The next set of challenges are concerned with the text mining approaches themselves. Most approaches depend on annotated corpora and sizable corpora based on full-text articles or other text sources we discussed are lacking. The largest full-text corpus, CRAFT (Bada *et al.*, 2012), contains 67 articles and the annotation focuses mostly on low-level semantic information, such as named entities and concepts. Some tasks we discussed require higher level annotation, such as annotation of argumentation, discourse structure, citation function and quotation, and are much more challenging since they are less well-defined and some subjectivity is involved in annotating them. Collaborative, cross-institution efforts would be beneficial for consolidating existing research in these areas and proposing more comprehensive characterizations. Ontology development research should also be taken into account, since some existing ontologies focus on scholarly discourse (e.g., SWAN (Ciccarese *et al.*, 2008)), and annotation efforts would benefit from the insights of such research. Another promising avenue is crowdsourcing of annotation, where the “crowd” (a mix of lay people, enthusiasts, experts), recruited through an open call, provide their services for a given task. In the biomedical domain, crowdsourcing has been successfully applied to relatively low-level tasks such as named entity annotation, while it has been considered less suitable for complex, knowledge-rich tasks (Khare *et al.*, 2015). However, the design of crowdsourcing experiments plays a significant role in their success and creative crowdsourcing interfaces could make collection of complex data (e.g., argumentation graphs) more feasible. It is also worth noting that frameworks like Nanopublications (Mons and Velterop, 2009) and Micropublications (Clark *et al.*, 2014) advocate the use of semantic models of scientific statements and argumentation, respectively, in the workflows of scientists as a means of knowledge generation and exchange. If such models are adopted more widely (not only among scientists but also publishers and other stakeholders), the knowledge generated would also be invaluable as gold standard data. The Resource Identification Initiative (Bandrowski *et al.*, 2016) promotes such a model for research resources (e.g., reagents, materials) and can be informative in this regard.

Representativeness and balance of a corpus is important for the generalizability of tools that are trained on it. Though corpus linguistics literature addresses the construction of balanced and representative corpora (e.g., (Biber, 1993)), in practice, most biomedical text corpora focus on a restricted domain of interest. For example, CRAFT (Bada *et al.*, 2012) contains biology and genetics articles, while GENIA (Kim *et al.*, 2003) contains abstracts about biological reactions involving transcription factors in human blood cells. Lippincott *et al.* (2011) showed that subdomains in biomedical literature vary along many linguistic dimensions, concluding that a text mining system performing well on one subdomain is not guaranteed to perform well on another. Construction of wide-coverage, representative biomedical full-text article corpora, while clearly very challenging, would be of immense value to text mining research in general. Also note that a subfield of machine learning, *domain adaptation*, specifically focuses on model generalizability. Various methods (some requiring data from the new domain and some not) have been proposed (e.g.,(Daumé, III, 2007)), and such methods have been applied to biomedical text mining tasks (e.g., (Miwa *et al.*, 2012a)). Independently, some systems provided machine learning models that can be retrained on new annotated corpora (e.g., (Björne and Salakoski, 2011; Leaman and Lu, 2016)), while others attempted to generalize by appealing to linguistic principles (e.g., (Rindflesch and Fiszman, 2003; Kilicoglu and Bergler, 2012)).

Important information in biomedical articles may only appear in tables, graphs, figures, or even supplementary files. There is relatively little research in incorporating data from such artifacts into text mining approaches, even though some semantic models, such as Micropublications (Clark *et al.*, 2014), support them. Figure retrieval has been considered, mainly focusing on using text from figure captions (Hearst *et al.*, 2007), text within figures (Rodriguez-Esteban and Iossifov, 2009), as well as text from paragraphs discussing the figures and NER (Demner-Fushman *et al.*, 2012). Research on information extraction from tables is rare (Wong *et al.*, 2009; Peng *et al.*, 2015; Milosevic *et al.*, 2016), though this may change with recent availability of corpora (Shmanina *et al.*, 2016). Jimeno-Yepes and Verspoor (2014) showed that most literature-curated mutation and genetic variant existed only as supplementary material and used open-source PDF conversion tools to extract text from supplementary files for text mining.

The accuracy of text mining approaches vary widely depending on the task. In some classification tasks (e.g., identifying PICO categories), state-of-the-art performance is over 0.9 accuracy, whereas in recognition of citation quotation, the state-of-the-art performance is just over 0.3. Although text mining tools have shown benefits in curation and annotation (Alex *et al.*, 2008; Névéol *et al.*, 2011b), it is critical to educate the users about the role of such tools in their workflows and their value/limitations, and not alienate them by setting their expectations impossibly high. It is also worth pointing out that human agreement on some tasks is not high; therefore, it may be unrealistic to expect that automated tools do well (e.g., Fleiss’ of 0.11 for deceptive text annotation (Ott *et al.*, 2011)). Depending on the task, a user may prefer not the setting which yields the highest F_1_ score, generally considered the primary performance metric, but rather high recall or high precision. Providing the ability to tune a system for high recall or precision is likely to be advantageous for its adoption. Most machine learning systems are essentially black-boxes, and the ability of systems to provide human-interpretable explanations for their predictions may also affect their adoption. Curation cycles, in which experts or the crowd manually “correct” text mining results, providing feedback that is automatically incorporated into machine learning models, can also be effective in incrementally improving performance of such models.

## 4 Conclusion

Towards enhancing rigor and integrity of biomedical research, we proposed text mining as complementary to efforts focusing on standardization and guideline development. We identified four main areas (plagiarism/fraud detection, compliance with reporting guidelines, management of information overload, and accurate citation), where text mining techniques can play a significant role and surveyed the state-of-the-art for these tasks. Among the tasks we discussed, we believe that the following can have the biggest and most immediate impact: given a document (e.g., manuscript, publication), i) checking for adherence to *all* elements of the relevant reporting guideline, ii) generating document-level and literature-level argumentation graphs, iii) constructing citation quotation networks. For some tasks, current state-of-the-art text mining techniques can be considered mature (e.g., extrinsic plagiarism detection, extracting PICO sentences); while for other tasks, substantial research progress is needed for practical tools (e.g., construction of argumentation graphs, identifying citation quotations). We argued that the main advantage of text mining comes in its ability to facilitate performing tasks at a large scale. By shortening the time it takes to perform tasks needed to ensure rigor and integrity, text mining technologies can promote better research practices, ultimately reducing waste and increasing value.

## Footnotes

1 https://grants.nih.gov/reproducibility/index.htm

2 https://grants.nih.gov/grants/research_integrity/whatis.htm

3 http://headt.eu

4 http://www.turnitin.com

5 http://www.plagramme.com

6 http://www.crossref.org/crosscheck

7 http://pan.webis.de

8 http://www.peneloperesearch.com/

9 http://www.statreviewer.com/

10 A service similar to Crossref’s CrossMark, which indicates updates on a given publication, can be envisioned.

11 Interesting from a research integrity/transparency perspective, the system developed in (Kilicoglu and Bergler, 2008) was used to compare the language used in reporting industry-sponsored research and non-industry-reported research, which found that the former was on average less speculative (ter Riet *et al.*, 2013).

12 http://www.nist.gov/tac/2014/BiomedSumm

13 https://report.nih.gov/

## Acknowledgements

I thank Jodi Schneider, Gerben ter Riet, Dina Demner-Fushman, Catherine Blake, Thomas C. Rindflesch, Olivier Bodenreider, and Caroline Zeiss for their comments on earlier drafts of this paper.

### Funding

This work was supported by the intramural research program at the U.S. National Library of Medicine, National Institutes of Health.

